# A novel weight lifting task for investigating effort and persistence in rats

**DOI:** 10.1101/752410

**Authors:** Blake Porter, Kristin L. Hillman

**Author notes:** Correspondence: Blake Porter.

## Abstract

Here we present a novel effort-based task for laboratory rats: the weight lifting task (WLT). Studies of effort expenditure in rodents have typically involved climbing barriers within T-mazes or operant lever pressing paradigms. These task designs have been successful for neuropharmacological and neurophysiological investigations, but both tasks involve simple action patterns prone to automatization. Furthermore, high climbing barriers present risk of injury to animals and/or tethered recording equipment. In the WLT, a rat is placed in a large rectangular arena and tasked with pulling a rope 30 cm to trigger food delivery at a nearby spout; weights can be added to the rope in 45 g increments to increase the intensity of effort. As compared to lever pressing and barrier jumping, 30 cm of rope pulling is a multi-step action sequence requiring sustained effort. The actions are carried out on the single plane of the arena floor, making it safer for the animal and more suitable for tethered equipment and video tracking. A microcontroller and associated sensors enable precise timestamping of specific behaviors to synchronize with electrophysiological recordings. The rope and reward spout are spatially segregated to allow for spatial discrimination of the effort zone and the reward zone. We validated the task across five cohorts of rats (total n=35) and report consistent behavioral metrics. The WLT is well-suited for neuropharmacological and/or *in vivo* neurophysiological investigations surrounding effortful behaviors, particularly when wanting to probe different aspects of effort expenditure (intensity vs. duration).

## 1 Introduction

Physical effort is often required to perform activities and reach goals. Subjects vary naturally in their willingness and ability to expend effort, with significant alterations in effort-based decision-making being a clinical feature of certain neuropsychiatric conditions (e.g., depression (Treadway et al. 2012, Yang et al. 2014)). To decipher the underlying brain mechanisms governing effort exertion (and dysfunctions therein), researchers need laboratory tasks that require physical exertion but that are also amendable to simultaneous neuroimaging, neurophysiological, or optogenetic techniques.

In rodent research, effort has generally been assessed using climbing barriers or operant lever pressing paradigms. The barrier-climbing paradigm, originally devised by Salamone et al. (1994), involves placing a vertical climbing barrier within a T-maze arm such that an animal must climb or jump – i.e., they must exert an extra degree of physical effort – to reach a reward site. The intensity of the effort can be increased by increasing the height of the barrier with 25-30cm being the most common. In rats, barrier paradigms have been used in lesion/inactivation studies (Walton et al. 2002, Rudebeck et al. 2006, Floresco and Ghods-Sharifi 2007, Holec et al. 2014, Karimi et al. 2017), pharmacological investigations (Schweimer and Hauber 2006, Bardgett et al. 2009), and electrophysiological recordings (Hillman and Bilkey 2010, Cowen et al. 2012) to assess the contribution of different brain areas and neurochemical systems to decisions which require physical effort. However, the protocol has limitations. Surmounting the barrier can become a simple, quickly executed motor action (i.e., a jump), especially when the barriers are small and/or the animal is frequently exposed to the apparatus. In theory effort difficulty can be increased, to an extent, by increasing the height of the barriers, however in practice this increases the risk of injury to the animal and/or tethered research equipment. Jumping into 3-dimensional space also complicates spatial tracking via an overhead camera and can generate noise in electrophysiological recordings.

In addition to barrier-climbing experiments, effort expenditure has also been investigated in rodents using operant lever pressing paradigms. Here, higher numbers of lever presses are equated with higher effort expenditure. Fixed ratio (FR) and progressive ratio (PROG) response schedules have been used effectively to probe the neurological mechanisms of effort-related cost-benefit decision-making (e.g., Floresco et al. 2008, Randall et al. 2014, Hart et al. 2017). The concurrent lever-press/reward choice paradigm in particular has been used to examine effort expenditure in relation to generalized behavioral activation (Salamone et al. 2002, Schweimer and Hauber 2005, Randall et al. 2012), with subtle pharmacological shifts in behaviors being produced by various compounds (see Salamone et al. (2018) for recent review). While lever pressing is an action that can be carried out alongside tethered optogenetic or electrophysiological experimentation (e.g., Ma et al. 2014, Robinson et al. 2014, Proulx et al. 2018, Lindenbach et al. 2019), lever pressing – even more so than barrier jumping – is a simple, quickly executed motor action. Hence the intensity of effort in FR and PROG lever pressing tasks is largely related to the repetition of responses over time, which introduces a temporal cost confound to effort costs when interpreting resultant data.

Directly increasing the intensity/difficulty of physical effort associated with a single lever press would better isolate an effort cost component. Holec et al. (2014) tested this idea by engineering weight-adjustable seesaw levers within the choice arms of a Y-maze. Lever weight was modulated as a percentage of each animal’s body weight, and the weight of a lever could be kept static during a single session or incrementally changed across trial blocks. While a novel paradigm, behavioral shortcomings were described in the report, including ceiling effects and failure to achieve pre-training criterion in a substantial number of subjects (Holec et al. 2014).

Due to the limitations of existing barrier and lever pressing paradigms, we aimed to design a task that: 1) was suitable for use with tethered cables and overhead tracking systems; 2) allowed the intensity of physical exertion to be directly modulated; and 3) involved an action that produced more noticeable/observable physical exertion – i.e., a more complex action sequence requiring sustained effort. Using Sprague-Dawley rats as subjects, we developed the Weight Lifting Task (WLT).

The WLT allows for behavioral characterizations of effort expenditure in laboratory rats, including those that are tethered for neurophysiological recording and/or optogenetic stimulation. In the WLT, the animal is placed in a large rectangular arena and tasked with pulling a rope 30 cm out of a rope conduit to trigger food delivery at a nearby reward spout; weights can be added to the rope in 45 g increments to increase the intensity of effort. As compared to lever pressing and barrier jumping, weighted rope pulling is a multi-step action sequence requiring sustained exertion. The actions are carried out on the single spatial plane of the arena floor, making it safer for the animal and more suitable for tethered equipment and video tracking. Automation of the WLT via an Arduino microcontroller enables precise timestamping of task components, which can be synchronized alongside neurophysiological recordings or stimulation. Thus the WLT is well-suited for neuropharmacological, neurophysiological, or optogenetic investigations of effort, particularly when different domains of effort are of interest (e.g., high-intensity exertion versus sustained persistence).

## 2 Materials and Equipment

### 2.1 WLT Arena

The arena is a wooden rectangle measuring 120 × 90 × 60 cm with all surfaces painted matte black. At the center of one wall is the rope conduit – a polyvinyl chloride (PVC) tube that extends 7 cm into the arena, and is elevated 1 cm above the floor (Figure 1). This rope conduit is used to guide the rope attached to the weight system into the arena. Aligned with the conduit on the arena floor is a 7 × 30 cm section of ribbed rubber to provide grip for the rat’s feet when pulling. Four cm left of the conduit a white light emitting diode (LED) is recessed into the arena wall to signal reward delivery. Seven cm left of the LED is the reward spout – a silicone tube (2.5 mm ID, 4.7 mm OD) that extends 20 cm into the arena at an approximate 45° angle to the wall and away from the rope conduit. This tube is used to deliver sucrose reward via a peristaltic pump; the pump is located outside the arena. The silicone tube is protected by an outer PVC tube (20 cm long, 3 cm in diameter) to prevent rats from chewing on the silicone tubing. At the end of the silicone tube spout is a plastic dish (3.5 cm diameter, 0.5 cm tall) to collect the sucrose. The rope PVC conduit and the silicone tube spout were, in later iterations of the task, separated by a wall which was 40 cm long, 20 cm high, and 4 cm thick (see *Discussion*). In the training phase (described below in *Methods*), a second, larger PVC tube measuring 25 cm long with a 3 cm diameter with horizontal slits down the sides is also inserted into the arena 12 cm to the right of the rope conduit.

**Figure 1:**
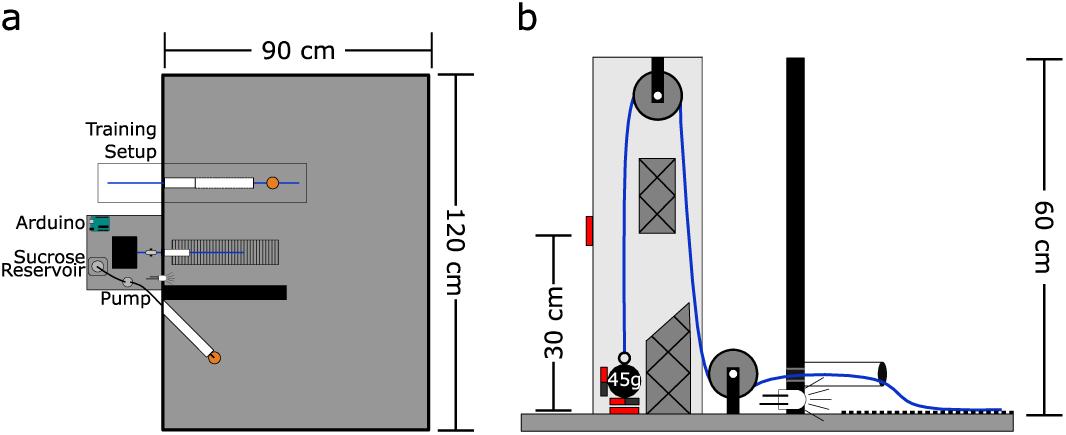
WLT schematic. a) Aerial view of task set-up. Outside the task area is the pulley system, Arduino microcontroller, and sucrose pump. Inside the task area, the training setup is outlined with the training tube (white) and training rope (blue) with sucrose dish attached (orange). Medial to the training setup is the working area containing the weight lifting rope conduit (white), weighted rope (blue), and rubber grip mat. The black wall divides the working area from the reward spout (white) and dish (orange). b) Profile view of the pulley system. Solid red boxes indicate magnetic reed switch placement. Styrofoam guide inserts are shown with cross hatches.

### 2.2 WLT Rope System

The rope system is comprised of a rope, two pulleys, and various weights (see Figure 1b). The rope is made of braided nylon and measures 4 mm in diameter with a length of 145 cm; one end of the rope extends into the arena for the rat to pull while the other end can be attached to a weight outside the arena. The rope runs through two nylon pulleys each with an outer diameter of 25 mm and track width of 8 mm (Zenith Inc.). The weight end of the rope uses a key chain clip to facilitate switching weights quickly. Lead fishing weights (bank sinkers; Maxistrike Inc.) are used to add weight as desired. Fishing weights were modified to range from 45 g to 225 g in 45 g increments. Each weight has 2-3 neodymium magnets (15 mm diameter × 4 mm thick) attached to it with two-part epoxy resin. The pulley set-up is enclosed in a wooden, open-faced box (7 × 9 × 60 cm tall) mounted so that the open face is oriented towards the arena. Two normally-open magnetic reed switches (Jaycar Electronics, Inc.) are embedded into the wooden housing, one switch is at the base where the weight statically sits and the second switch is 30 cm above the base. Two Styrofoam inserts within the wooden housing help to prevent the weight from swinging, and to keep the weight close to the reed switches to ensure they are triggered.

### 2.3 WLT Automation

An Arduino Uno microcontroller (www.arduino.cc) is used to control the experiment. The two magnetic reed switches feed into the Arduino which controls the LED and peristaltic pump (12 Volt; Adafruit Industries, LLC) for sucrose delivery. Adafruit’s “Motor Shield V2” for Arduino is used to power and control the peristaltic pump. The Arduino and pump are run off of a 12 Volt, 4.5 Amp hour lead-acid battery (DiaMec Limited) to reduce electrical line noise during electrophysiology experiments. Rats have to pull the rope 30 cm in order to trigger the reed switch located 30 cm above the pulley system base. If this switch is triggered, the Arduino turns on the LED for 250 ms and 0.2 mL of 20% sucrose solution is dispensed through the peristaltic pump. In the rare instance where a rat makes a successful pull and the reed switch fails to trigger, the Arduino has a button wired to it for manual dispensing of sucrose and LED illumination. This button also aids in autoshaping the rats during training (see Methods below). The Arduino is configured to send TTL signals to a Neuralynx acquisition system (Digital Lynx SX; Neuralynx Inc), such that all weight pulling events can be timestamped alongside neural recordings and video tracking. The Arduino signals: when the weight first leaves the base reed switch; when the weight reaches the 30 cm reed switch (or the experimenter uses the button); and when the weight returns to the base reed switch. This allows for capture of both successful pulls (rats pulling up the weight a full 30 cm for a reward) and unsuccessful pulls (lifting the weight but failing to lift it to 30 cm). A capacitive touch lick sensor can also be added to the reward dish. However, we found that this causes electrical noise when performing *in-vivo* electrophysiological recordings so we did not continue with this sensor feature. The Arduino code for the WLT is available on Github (https://github.com/blakeporterneuro/weightLiftingTask).

## 3 Methods

### 3.1 Subjects

Thirty-five male Sprague-Dawley rats (450-650 g) were used in total to validate the WLT. These were run as five separate cohorts (7 + 6 + 8 + 4 + 10 rats) by two different experimenters over an 18 month timespan. All rats were 2-6 months old at the start of the experiment and obtained from the University of Otago’s Hercus-Taieri Resource Unit. Rats were housed in groups of two in plastic individually-ventilated cages (38 × 30 × 35 cm). The animal housing room was maintained on a 12 hour reverse light-dark cycle and kept between 20 – 22°C. Rats were given two weeks from the time of arrival to acclimate to the new facility. During this time rats had *ad libitum* access to food (18% Protein Rodent Diet; Tekland Global) and water. After the acclimation period, each rat’s free-feed weight was measured and rats were food deprived to no less than 85% of their free-feed weight throughout the experiment. Rats always had *ad libitum* access to water. All experiments were carried out during the dark phase. All experimental protocols were approved by the University of Otago Animal Ethics Committee and conducted in accordance with New Zealand animal welfare legislation.

### 3.2 Habituation and Autoshaping

For three days rats were habituated to the experimental room and the experimenter by being handled on the experimenter’s lap for 10 min/day. Starting on day four, rats spent one min in the experimenter’s lap before being placed in the arena for 10 min/day. On days four through six, small drops of 20% sucrose solution were randomly scattered around the arena to promote interest in this food reward.

From day four to approximately day 22, rats were autoshaped to pull a rope for a sucrose reward. This was initially achieved by placing a “training rope” completely inside the arena. The training rope was 60 cm long and had a sucrose reward dish – identical to the dish located at the usual reward spout – epoxied at its midpoint. Sucrose could thus be obtained in the dish on the training rope (“training dish”, ∼0.2-0.5 mL) and/or at the usual reward spout (“reward dish”). Each time a rat consumed sucrose from the training dish, sucrose was also dispensed to the reward dish via a button press to the Arduino. Sucrose was replenished in the training dish by the experimenter using a syringe when the rat was at the reward dish. Once rats were readily consuming sucrose from both the training and reward dishes, one end of the training rope and the training dish were inserted into the PVC “training tube” (see *Materials and Equipment* and Figure 1a). The other end of the training rope was extended out of the arena so that the experimenter could manipulate the position of the training dish.

Initially, the training dish was only partially inserted into the training tube, such that it would be easily accessible for the rat the reach in and retrieve the dish. Rats would generally pull the training dish out of the tube with their teeth or forelimbs. As rats became more familiar with this procedure, the training dish was put further and further into the tube – away from the arena opening – after each training sucrose consumption. The critical part of autoshaping occurred when the training dish was too far inside the training tube to grab directly and rats needed to pull the training rope to retrieve the dish. In our experience, some rats would lose interest in the training dish when it was no longer within reach of teeth or forelimbs. When this occurred, the training dish was placed closer to the arena opening so that the rat could once again retrieve the sucrose. Once consumption behavior was reinstated, the process of incrementally putting the training dish further and further into the tube – away from the arena opening – was repeated. The maximum distance that the training dish was placed inside of the training tube was 12 cm from the arena opening.

### 3.3 WLT Training

Once the rats learned to consistently pull the training rope to get the training dish out of the training tube, WLT training commenced. Now, sucrose was no longer provided in the training dish and was only provided via the usual reward spout. Rats were first placed in the arena and allowed to perform 5 trials using the training rope extending from the training tube. The training rope and tube were then removed, leaving only the regular WLT rope conduit and reward spout (see Figure 1a). Rats would then learn to pull the regular, non-training rope for sucrose reward; the rope was not weighted with lead weights at this stage (“0 g”) but did carry the weight of the magnets (∼5 g). Initially, rats would be manually rewarded (via the Arduino button) for very small pulls on the regular rope. As training progressed, rats would need to pull the rope further and further to get rewarded. Rats were trained on this “0 g” level (no lead weight, only magnets) until they were making successful 30 cm pulls to trigger automated reward delivery on greater than 80% of their attempted pulls. After this, training sessions consisted of 10 successful 0 g pulls followed by addition of a 45 g lead weight (“45 g”). WLT training was deemed complete when rats were able to successfully pull the 45 g weight on more than 80% of attempts and completed 10 successful attempts each of 0 g and 45 g in less than five min.

### 3.4 Surgical Window

After reaching the WLT training criterion, rats underwent surgical implantation of electrode arrays. This was a one day surgery, involving stereotaxic craniotomies under isoflurane anesthesia, as previously described (Porter et al. 2019). Rats were given 10 days of post-operative recovery and then re-tested on the WLT using the last training parameter, i.e., 10 successful attempts each of 0 g and 45 g in less than five min. Rats were now performing the WLT with a headplug connected to a headstage (Neuralynx HS-36-LED or HS-32-mux-LED), tethered to a commutator (3 meter tether, Neuralynx Saturn-1). All rats achieved the WLT training criterion within one to eight days of re-testing. Electrophysiological data are not analyzed in this manuscript, however we mention this surgical window here to demonstrate that the WLT is conducive for use in surgically implanted, tethered animals. Example LFP traces from the anterior cingulate cortex of a rat making 10 successful pulls on 135 g can be seen in Supplemental Figure 1.

### 3.5 Behavioral Experiments – General Design

Each experiment described below was performed in this general sequence: two min pre-baseline, experimental task, two min post-task baseline, satiation check. In the two min pre-baseline, the animal was placed in the arena but the rope was not available. The purpose of this pre-baseline was to collect two min of neural and locomotor behavior when not performing the task. At the two min mark the rope was inserted through the rope conduit and the experimental task was immediately started. At the end of the task, the rope was again made unavailable and the rat remained in the arena for two min to enable collection of end-of-task neural and locomotor behavior. Two sucrose rewards were then manually delivered via the reward spout. The purpose of this was to determine if the rat was still motivated to consume sucrose, or satiated. Ready consumption of two rewards was scored as “non-satiated”, one of two rewards as “partially satiated”, and none of the rewards as “satiated.” The two min pre- and post-recordings successfully provided non-task, ‘open field’ behavior as compared to the experimental task period (Figure 2). For the experimental task period, we defined two spatial regions of interest (ROIs) for subsequent analysis purposes: the on-task ROI and the off-task ROI. The on-task ROI was defined as the 45 × 50 cm area encompassing the rope conduit and reward spout. Rats also had to be making attempts while in the on-task ROI in order to be considered on-task. The off-task ROI was designated as the remaining area of the arena outside the on-task ROI as well as when the rats were in the on-task ROI but making no attempts at pulling the rope.

**Figure 2:**
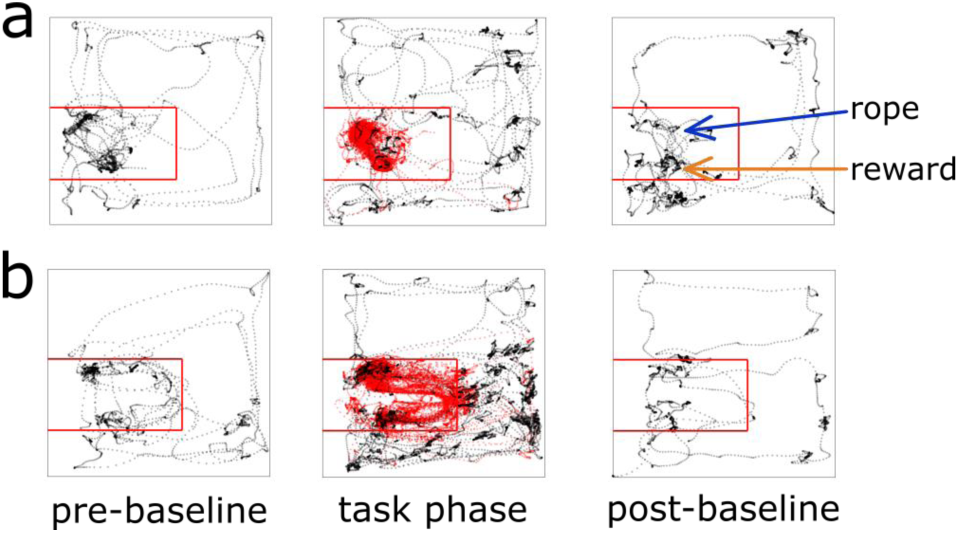
Animal tracking and ROI examples. Tracking data from two different recording sessions is shown; panel (a) is a session from an early iteration of the task where there is no wall between the rope and reward, panel (b) illustrates a session where the wall is present. The on-task ROI is outlined with the red square. Rat tracking data is shown as black dots if off-task and red dots if on-task.

#### 3.5.1 Progressive Weight Paradigm

The progressive weight paradigm used progressively heavier weights to increase effort intensity across time. After the two min pre-baseline, the weight rope with 0 g was inserted into the arena. After 10 successful trials, 45 g was added to the end of the rope. This was repeated every 10 successful trials until either 225 g was reached or the rats quit the task. Quitting was defined as the rats making no attempts to pull the rope for one min (cohorts 1-2) or two min (cohorts 3-5; empirically we had determined from the initial cohorts that one min was too short of a duration to define quitting). Although rare, if a rat managed 10 successful trials on 225 g, the task would be made “impossible” by wrapping the rope around a solid bar outside the arena to prevent it from being pulled high enough to trigger a reward. We refer to the amount of weight at this stage as “infinity”. Rats would never receive a reward during this impossible phase despite their persistent, frustrated efforts. However, rats often quit before completing 10 successful 225 g trials.

#### 3.5.2 Fixed Weight Paradigm

The fixed weight paradigm used a fixed weight of 180g to investigate persistence and quitting in a fixed difficulty context. The fixed weight was determined for each rat based on their performance on the progressive weight paradigm – their highest achievement weight was used, that is, the highest weight on which the rat completed 10 successful trials. For most rats the fixed weight was 180 g fter the two min pre-baseline, the weight rope with 0 g was inserted into the arena. After 10 successful trials, the fixed weight was immediately added to the end of the rope. Rats could complete as many trials as desired until they quit or until one hour elapsed, whichever came first. Quitting was defined as the rats making no attempts to pull the rope for two minutes.

### 3.6 Data Analysis

All data analyses were carried out using custom Matlab scripts. First, Neuralynx TTL events and tracking data were imported into Matlab along with an info txt file that contained the times the weights were changed (e.g., when 45 g replaced 0 g) and when the rat quit. Time spent on each weight was calculated by the duration it took the rat to complete 10 trials of a weight or, for the quit weight, the duration from when the weight was attached until the rat quit. The duration of a trial was calculated by the time between reward TTL signals. The number of attempts the rats made for each weight was determined by the number of times the weight was lifted high enough to trigger the reed switch at the base of the weight lifting apparatus. Attempts were further broken down into successful and failed attempts. Successful attempts were attempts where the rat pulled the weight high enough to trigger a reward. Failed attempts were attempts where the rat lifted the weight but not high enough to trigger a reward. The quit weight was the weight in which the rat did not complete 10 trials and stopped making attempts for two minutes. The achievement weight was the highest weight the rat completed 10 successful trials on. Time on-task was determined by calculating the time that the rat was located in the on-task ROI of the arena and was making attempts while within this ROI. If the rat left the on-task ROI but returned within three seconds he was still considered on task. If the rat was not present in the on-task ROI or in the on-task ROI but not making any attempts while in the ROI, they were considered to be off-task. In order to analyze the fixed weight paradigm over time, we took the first, middle, and last 30 trials on 180g when analyzing the percentage of failed attempts. For analyzing the duration of successful trials we took the first, middle, and last 10 successful trials on 180g. All data were first tested for normality using D’Agostino & Pearson normality test before the appropriate statistical test was conducted.

## 4 Results

### 4.1 Shaping and Training of Weight Pulling

After three days of habituation to the apparatus, rats began the shaping procedure using a training rope outfitted with a sucrose training dish (see *Materials and Equipment, Methods*). Shaping stages were defined as: i) consuming sucrose readily from both the training sucrose dish and reward spout; ii) retrieving the sucrose dish readily with forelimbs/teeth when the training dish is placed progressively further inside of the training tube (Figure 3a); iii) retrieving the training dish readily by pulling the attached training rope when the dish could no longer be reached inside the training tube (Figure 3b, top); iv) pulling the training rope with no sucrose in the training dish and receiving sucrose only from reward spout; v) transitioning from the training rope to the 0 g weighted rope (Figure 3b, bottom); and vi) reaching WLT training criteria of 10 trials each of 0 g and 45 g within five min. Shaping stages i to iv can be seen in Supplemental Video 1.

**Figure 3:**
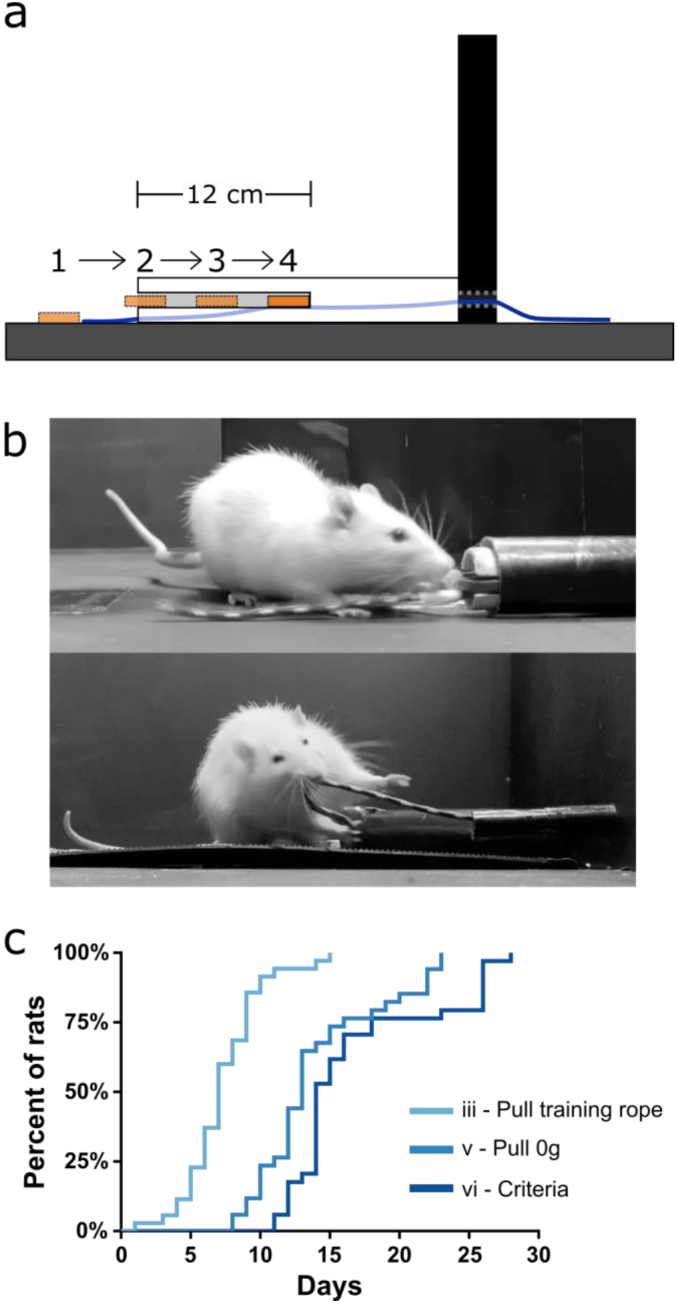
WLT training. a) Schematic showing profile of training tube and the progressive placement of the sucrose dish further and further inside the training tube. b) Picture of a rat learning to pull the training dish (stage ii; top) and pulling the 0 g weighted rope (stage v; bottom) c) Training data for 35 rats indicating mastery of stages iii, v, and vi.

After the initial three days of habituation, it took 7.3 ± 2.8 days (mean ± SD) for rats to reach stage iii and become proficient in pulling the training rope (Figure 3c). Transitioning from the training rope to the WLT rope with 0 g (stage v) took an average of 14.0 ± 4.5 days while reaching stage vi proficiency required an average of 16.9 ± 5.4 training days. We found that training frequency was an important consideration. Anecdotally, conducting shaping and training seven days/week tended to be more successful than taking weekend breaks, where rats would regress a stage or two after each two day break.

The most critical and arduous step in shaping was the transition from stage ii to stage iii, where the training dish was out of forelimb/teeth reach in the training conduit. At this stage rats had to learn to pull the rope rather than the dish. Initially, rats become quickly uninterested in the dish when it was out of reach. This was remedied by moving the dish back within reach to reinstate interest in the sucrose reward (see *Methods*). An additional strategy to aid in the stage ii to iii transition was to initially place the training dish within a rat’s reach within the training tube but then as the rat approached, the experimenter would pull the dish (via the end of rope outside the arena) such that the dish was no longer within reach of the rat. This encouraged the rats to scramble with their paws for the dish and happen upon pulling the rope (see Supplemental Video 1 at 0:54 seconds). Out of 35 rats trained on the WLT, one rat never overcame this within-reach/out-of-reach obstacle despite lengthy shaping sessions (more than 30 days) and was removed from further study. Thus in our experience, the WLT shaping period is relatively short and has a high success rate, with 97% of our subjects reaching training criteria in under four weeks (Figure 3c).

In our experience, all rats developed the strategy of grabbing the rope in their teeth, pulling with their bodies, then holding the rope in their forepaws before pulling again with their teeth (see Supplemental Video 2). Some rats would, on low weights (0 and 45 g), simply hold the rope in their teeth and run away from the conduit until the reward triggered. However, this running strategy was not feasible for heavier weights and generally extinguished over time. To facilitate uniform pulling behavior and consistent effort loads during the shaping and training phase, if rats tried to pull the rope out of the conduit at 90° angles to the conduit, the experimenter held the rope before it reached the reward trigger height to discourage this behavior.

### 4.2 Progressive Weight Paradigm

In this experiment rats were tasked with progressively heavier weights after every 10 successful trials. The experimental session started with 0 g and the weight was increased in 45 g increments until either the rat quit the task or a pulling weight of 225 g was reached. The weight of the rope at time of quitting was deemed the “breakweight” in line with PROG-lever pressing “breakpoint” terminology. Of the 35 rats trained on the task, 22 of them have completed the progressive weight paradigm across 157 sessions (average of 7 ± 1 SD sessions per rat); behavioral data are shown in Figure 4. The most frequent breakweight observed was 225 g, occurring on 54 out of the 157 sessions (34%; Figure 4a). Of the 157 sessions there were only seven sessions where a rat achieved 10 successful trials on the 225 g weight and progressed to the “infinity” stage described in the Methods. Thus “infinity” data is provided in Figure 4a but is absent from other panels as there were so few occurrences and rewards (successes) never occurred on this condition.

**Figure 4:**
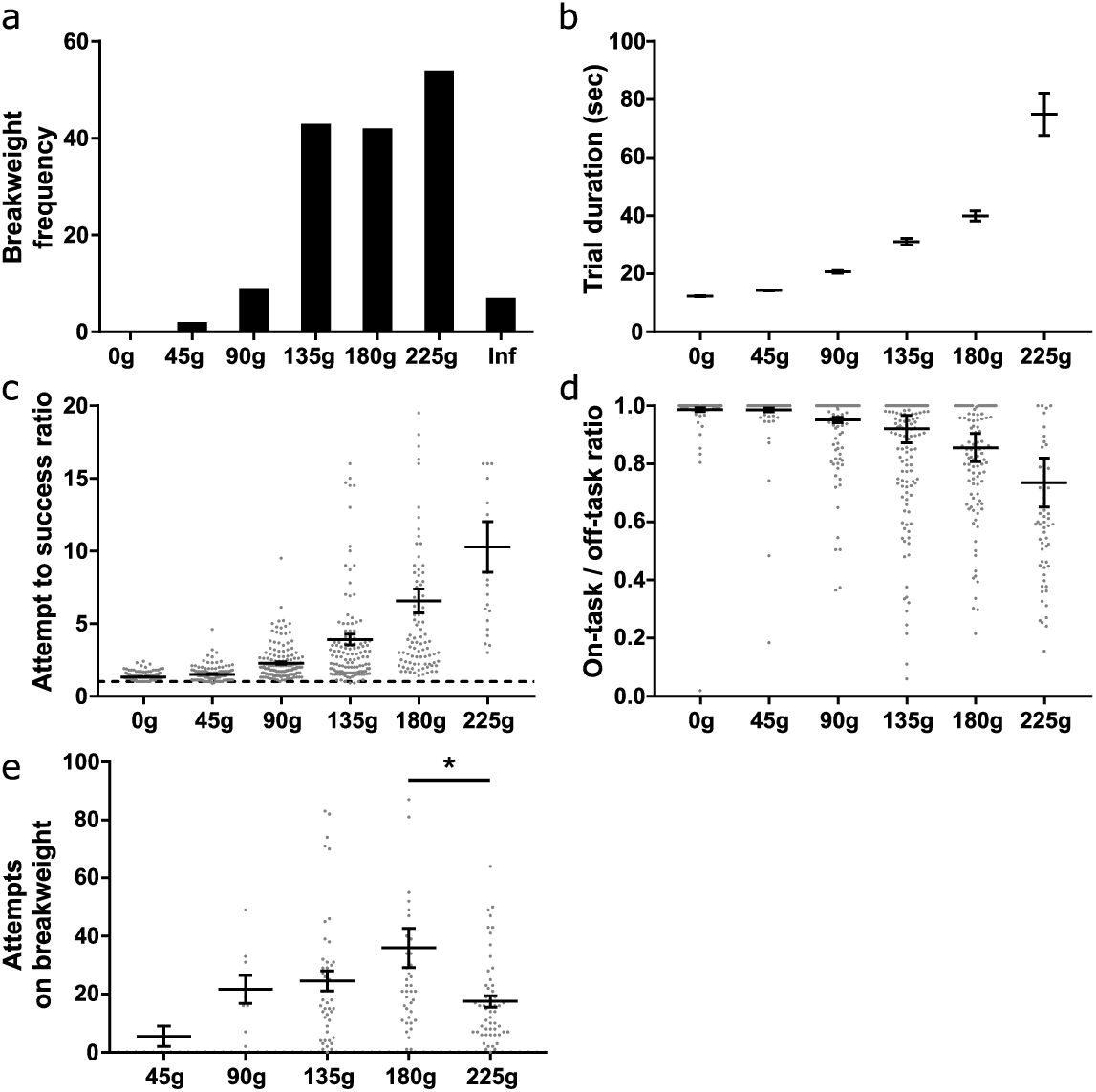
Progressive weight behavioral analyses. a) Breakweight distribution across all sessions. b) Average trial durations for each weight as measured by the time between successful trials. c) The ratio of attempts to successful trials for each weight. Dashed line indicates a 1-to-1 ratio. d) Ratio of time spent on-task / off-task for each weight. e) The number of attempts rats made on a session’s breakweight before quitting. Throughout the figure, grey dots indicate individual sessions, bars indicate mean ± 1 SEM.

The progressive weight paradigm exhibited predictable relationships between behavioral metrics associated with increasing effort and increasing weight. As the weight got heavier, trial duration significantly increased (KW (6) = 2277, p < 0.0001; Figure 4b) likely due to the rats failing more often in their attempts to pull the rope the full 30 cm (KW (6) = 371.5, p < 0.0001; Figure 4c). Furthermore, as the weights got heavier rats spent more time off-task (KW (6) = 271, p < 0.0001; Figure 4d). Specific examination of the breakweight trial blocks revealed a significant main effect for the number of attempts made on the breakweight before quitting (KW (6) = 12.22, p = 0.032; Figure 4e). However, a great deal of variation can be seen for each breakweight where some rats make many attempts before quitting while others quit after just a few attempts. Satiation checks carried out after the quit point (see *Methods*) were always 100% successful, suggesting that animals had not quit the WLT due to sucrose satiation.

In order to get a better understanding of the rats’ quitting behavior we broke down breakweights by individual rats and by session day. There was a main effect for rat on average breakweight (KW (22) = 56.42, p < 0.0001) indicating that individual rats had different breakpoints (Figure 5a). This variance was unrelated to body size differences between individual rats, as animal weight and average breakweight was not correlated (R^2^ = 0.004. p = 0.79; Figure 5b). Breakweight was significantly influenced by session day (KW (8) = 21.4, p = 0.003; Figure 5c), largely driven by day 1 which had a significantly lower average breakweight compared to days 4, 6, 7, and 8 (all p’s < 0.05; Dunn’s test for multiple comparisons). No other pairwise comparisons were significantly different. Taken together, the progressive weight paradigm is suited for investigating the effects of incremental changes in effort on persistence behaviors and quitting behaviors.

**Figure 5:**
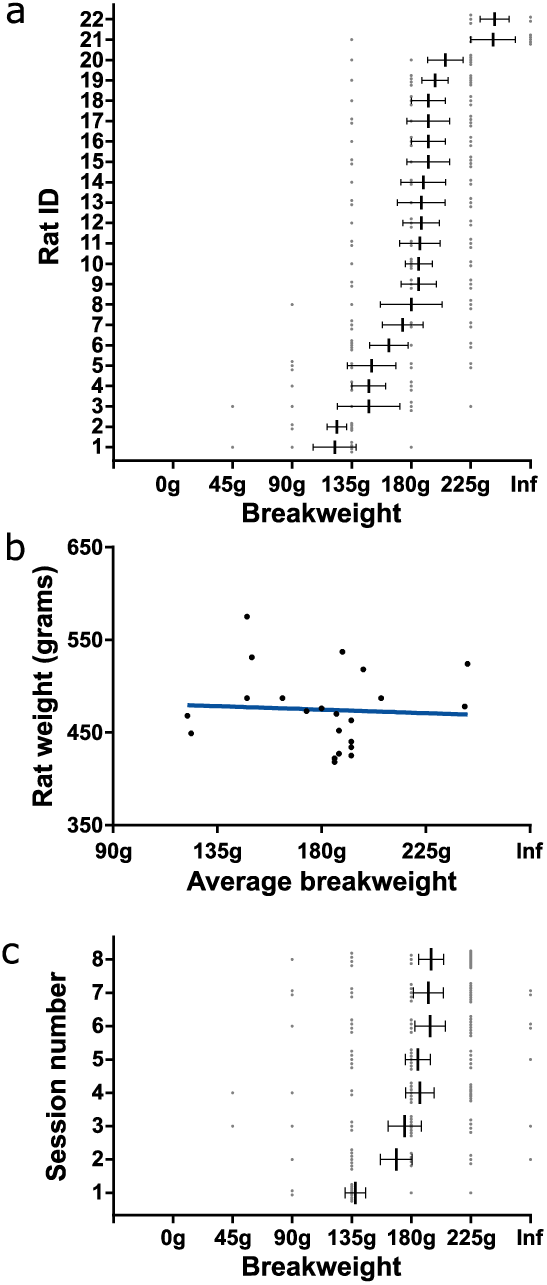
Detailed quitting behavior on progressive weight paradigm. a) Individual rat’s breakweight. b) Rat’s average breakweight by body weight. Blue line indicates line of best fit. c) Average breakweights across consecutive sessions. Throughout the figure, grey dots indicate individual sessions, bars indicate mean ± 1 SEM.

### 4.3 Fixed Weight Paradigm

In this experiment, rats were tasked with pulling a fixed weight (180 or 225 g) for as long as desired within a 60 min window; there were no progressive increases in weight. Ten trials on 0 g was used to start the session, after which the higher weight (180 or 225 g) was immediately attached. Eleven rats that carried out the progressive weight paradigm were subsequently tested on this fixed weight paradigm. Ten of these rats were tested with a 180 g fixed weight while one rat had 225 g. Across the 10 rats, 57 fixed weight sessions were completed in total, with each rat contributing three to six sessions. Performance on the fixed weight paradigm was variable across sessions and rats. Nonetheless, the number of attempts on the fixed weight before quitting, and the total time spent on the fixed weight task, both fit normal distributions (p = 0.93 and p = 0.67 respectively, D’Agostino & Pearson normality test; Figure 6a, 6b).

**Figure 6:**
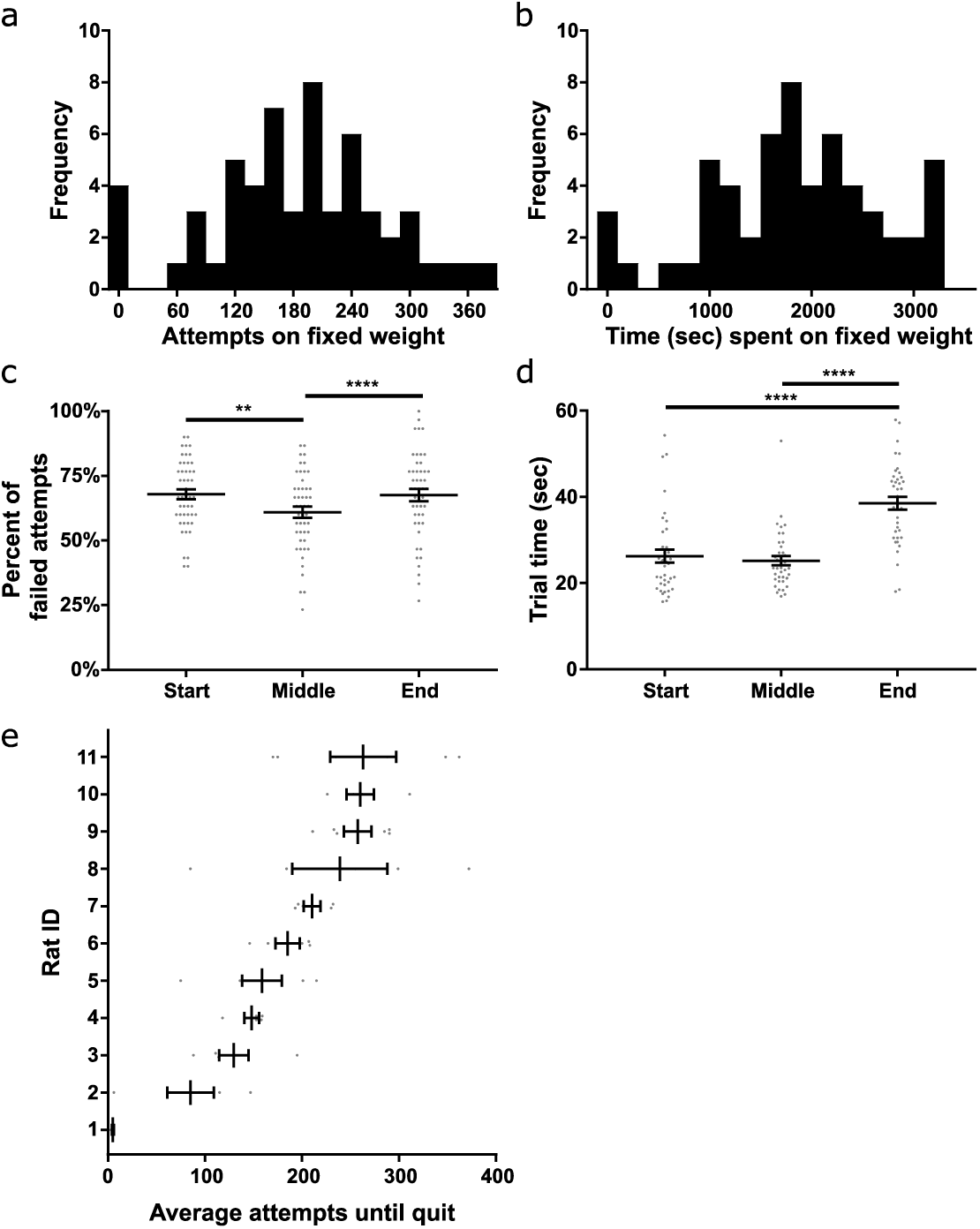
Fixed weight behavioral analyses. a) Histogram of the number of attempts made on the fixed weight (180 or 225 g) before quitting. b) Histogram of the time spent on the fixed weight before quitting. c) Percent of failed attempts and d) time for each successful trial at the start, middle, and end of the session. e) Individual rat’s average attempts on the fixed weight before quitting. Throughout the figure, grey dots indicate individual sessions, bars indicate mean ± 1 SEM.

We tested whether or not performance on the fixed weight changed over time, presumably due to fatigue developing across the session. Sucrose satiation checks (see *Methods*) were always 100% successful at the end of the task, suggesting that performance changes were likely unrelated to satiation. Time had a significant effect on the percent of failed attempts to all attempts (F (2) = 4.48, p < 0.0001, RM ANOVA; Figure 6c). Multiple comparisons testing revealed a significant difference between the failure ratio of pulls when comparing the start of the session to the middle of the session (p < 0.008), as well as when comparing the middle of the session to the end of the session (p < 0.0001; Holm-Sidak’s test). Anecdotally, rats tended to fail when the weight was immediately changed from 0 g to the heavier fixed weighted, after 10 successful pulls on 0 g. The rats would then acclimate to the heavier weight and the percent of failed pulls would reduce in the middle of the session, before increasing again towards the end of the session prior to quitting. Furthermore, time had a significant effect on the speed at which rats completed successful trials (F (3) = 34.67, p < 0.0001, Friedman’s test; Figure 6d). Rats slowed down significantly towards the end of the session – prior to quitting – as compared to the start of the session (p < 0.0001) and the middle of the session (p < 0.0001, Dunn’s test).

We further broke down fixed-weight task behavior by individual rat, and found a significant difference in the number of attempts before quitting across rats (KW (11) = 39.66, p < 0.0001; Figure 6e). Rat #1 in particular hardly performed the task over three days, generally making three successful attempts on the fixed weight and then quitting, despite doing 10 pulls of the same weight (180 g) only days prior on the progressive weight paradigm. The number of attempts made before quitting was not significantly correlated to rat body weight (all rats: R^2^ = 0.28, p = 0.10; excluding Rat #1: R^2^ = 0.25, p = 0.14). Overall, the majority of rats we tested were willing to perform the fixed weight paradigm for extended durations, making the task suitable for investigations of fatigue and persistence.

## 5 Discussion

Here we report a novel weight lifting task that can be used to investigate effort-based behaviors in rats. Rats can be trained on the WLT within a reasonable timeframe and are willing to carry out the positively-reinforced task. Once rats are trained on weighted rope pulling, the WLT can be used in a variety of ways to test different aspects of effortful behavior. We systematically tested two versions of the task – the progressive weight paradigm and the fixed weight paradigm – each modeled after traditional operant box PROG and FR response schedules. The progressive weight paradigm allows for investigating the role of increasing effort intensity on behavior. In contrast, the fixed weight paradigm is better suited for long term effort expenditure, endurance, and persistence. Many other experimental paradigms are possible – such as a choice-based decision-making WLT – due to the flexibility of the WLT. The WLT is constructed from inexpensive, easy to obtain components. Task automation and event detection via an Arduino allows for user-friendly, low cost implementation for labs looking to enhance their effort behavior investigations.

Holec and colleagues (2014) were the first, to our knowledge, to develop a rodent weight lifting-type task to investigate effort-based behaviors. They utilized weighted levers, one at the end of the two choice arms of a Y-maze, as a means of weight lifting. The weight required to depress the lever was chosen based on the animal’s body weight, with a maximum value of 40% of the rat’s body weight. Their weighted lever task was part of a task battery used to investigate the role of the anterior cingulate cortex (ACC) in effort behaviors and decision-making. In previous studies that have utilized climbing barrier tasks, lesioning or neurochemically manipulating the ACC has been shown to bias rats away from choosing effortful high-cost, high-reward (HCHR) choices and towards low-cost, low reward (LCLR) choices (Walton et al. 2002, Rudebeck et al. 2006, Schweimer and Hauber 2006). In contrast, Holec et al. found that ACC lesions did not have a large impact on rodent’s effort preference in the weighted lever task when using 20% of body weight. However, when Holec et al. repeated the experiment with a higher effort cost (40% of body weight), many behavioral issues were reported. For example, 8/20 rats could not complete the training phase of the task. Furthermore, behavioral results were difficult to interpret as four ACC lesioned rats showed no difference in HCHR preference as compared to controls, while the other two ACC lesioned rats essentially never chose the HCHR option. Their findings that ACC lesions may affect some effort behaviors (barrier jumping) but not others (20% value weight lifting), makes an important distinction in effort behavior research. We think our WLT – which requires more complex motor movements as compared to lever pressing, and fewer training and behavioral difficulties as compared to weighted lever pressing – could help investigators better elucidate subtle differences in effort exertion, such as those reported by Holec et al. (2014).

Our weight lifting task overcomes some of the common problems encountered in traditional effort-based tasks that use climbing barriers (Salamone et al., 1994) or operant box lever pressing (e.g., Floresco et al., 2008). While climbable barriers have been used successfully to investigate effort behaviors to date, climbable barriers have inherent experimental constraints. Experimenters can only make barriers so tall – and thus effortful – before rats either refuse to make attempts or do make an attempt but fail, resulting in the possibility of animal injury and/or damage to hardware devices. Our WLT allows for fine control over the amount of effort (weight amount) necessary to carry out the task. Furthermore, if a rat fails on lifting a weight there is no possibility of injury to the rat or damage to equipment, and any neurophysiological signals being recorded remain in-tact. While we have not carried it out, the WLT could be designed as a choice-based, decision-making task by putting a pulley system at the end of each arm of a Y-maze, similar to Holec et al.’s (2014) weighted lever task. Different weights or reward amounts could then be used to create traditional HCHR vs LCLR choice paradigms.

Additional paradigms could also be easily implemented using the WLT arena we have detailed here, that is, one with a single pulley system and an Arduino. For example, a progressive ratio schedule could be programmed into the Arduino requiring an increasing number of successful pulls to obtain a reward. Other weight and reward manipulations are also possible. For example, we have piloted a paradigm where, after a number of successful pulls on a low weight (e.g., 45 g), the task becomes impossible (“infinity weight,” see *Methods*) and no reward can be obtained. This paradigm lends itself well to effort-based reinforcement learning and investigations into frustration as rats become very annoyed when faced with the infinity weight situation.

Our WLT also confers benefits over operant box lever pressing effort tasks. Lever pressing tasks use the number of lever presses as the metric for effort. Effort-based lever pressing tasks generally use a fixed number of presses or a progressive ratio of increasing press numbers required to obtain a reward (e.g., Floresco et al. 2008, Randall et al. 2014, Hart et al. 2017). Number of lever presses has also been used as an effort metric in non-human primate effort studies (Kennerley et al. 2009). While operant box lever pressing tasks work well with tethered animals, the simple act of lever pressing does not lend itself well to study sustained, effortful action execution. Furthermore, using the number of lever presses to manipulate effort has a correlated confound of time making it difficult to parse behavioral changes due to the effort of many lever presses or due to the temporal discounting of rewards. Our WLT avoids this issue as the rats must always perform the same action (pulling the rope 30 cm) while the intensity of effort associated with that action can be manipulated via the attached weights. Furthermore, rope pulling is a more prolonged sequence of motor actions that may be better suited for studying the brain mechanisms behind effortful action planning and execution.

In addition to improving upon existing rodent-based effort tasks, we suggest that our rodent-based WLT offers a better behavioral comparison to the effort tasks used in non-human primate and human-based research. The primary motor-based effort task used with non-human primates and humans is grip-force (e.g., Pessiglione et al. 2007, Kurniawan et al. 2010, Varazzani et al. 2015). Generally, participants need to grip and squeeze a force meter with their dominant hand for a sustained time period or/and for a certain level of force. The grip force task is widely used as it can be done in a variety of experimental settings such as during EEG recording (Harris and Lim 2016) and fMRI scanning (Klein-Flugge et al. 2016). Our WLT is similar in nature as rats must pull the rope for a sustained period of time and with an appropriate level of force to obtain a reward. In contrast, barrier jumping or lever pressing is a single, quick exertion of effort. We hope that the WLT can be used with a variety of manipulations to help bridge the gap between human effort behavioral studies and rodent effort behavioral studies.

One limitation in early iterations of developing the WLT was the proximity of the rope to the reward spout. Rats figured out that they could pull the rope to the reward spout and get rewarded there with minimal movement between the rope area and reward area. To better spatially and temporally segregate the working area from the rewarded area, we placed a wall between the rope conduit and reward spout (see Figures 1 & 2, *Materials and Equipment*). This wall had the additional benefit of keeping the rats on the rubber mat. Without the wall, rats would sometimes try to pull the rope while standing on the wooden arena floor and this would result in the animals slipping, especially on weights above 90 g.

We specifically designed the rope conduit and reward spout to extend from the apparatus wall in order to prevent tethered rats from hitting their implants on the arena walls, which can produce electrophysiological artefacts. It would be feasible to outfit a bespoke operant box with the WLT for high throughput behavioral studies. However, in our experience, rats will need at least 35 cm of space in front of the rope in order to pull the rope successfully. In addition, we purposefully used a large arena because it allowed us to spatially segregate different behaviors. Anecdotally, when rats would grow frustrated with the task or when they would quit, they would sprint around the large arena then groom in a corner (see tracking data in Figure 2). Such nuanced behaviors may not be captured when using a more confined operant box.

We think it is important to discuss the behavioral variability produced by our WLT and the value of this variability. Performance across rats can be quite variable, and variability was also observed within a rat’s day-to-day performance. Figure 5 and Figure 6e depict this variability showing that some rats are willing to exert much more effort as compared to others. Furthermore, individual rats may, on some sessions, work very hard while on other sessions give up quickly. Overall, however, all but one rat we have tested was able to successfully pull 180 g (roughly 38% of average body weight, min: 31%, max: 43%). Thus, while there is rat-to-rat and day-to-day variability, all rats are able to carry out the task to a high degree of proficiency; comparisons across weights and across rats is feasible. Importantly, this variability in performance is not simply correlated with the rat’s body weight. We think this variability could lead to exciting investigations into the neural mechanisms underlying motivation, persistence, and quitting behaviors, including individualized intrinsic levels of motivation. In addition, the WLT is well-suited for the recent advances in animal behavioral tracking analyses such as DeepLabCut (Mathis et al. 2018) or DeepBehavior (Arac et al. 2019) which provide highly detailed, three dimensional kinematic data. For example, the motor action sequence of pulling the rope is quite complex compared to a lever press or jump, and likely requires extensive motor planning and sensory feedback for successful performance. The wide repertoire of behaviors elicited by the WLT, such as complex motor movements, reward consumption, task approach and avoidance, and quitting – when coupled with neurophysiological techniques – can provide a better understanding of the neural circuits involved in effort-based behaviors (Krakauer et al. 2017).

## 6 Conflict of Interest

*The authors declare that the research was conducted in the absence of any commercial or financial relationships that could be construed as a potential conflict of interest*.

## 7 Author Contributions

B.P. and K.H. designed the experiments. B.P. built the WLT, programmed the Arduino, trained the rats, and ran the experiments. B.P. analyzed the data and B.P. and K.H. interpreted the results. B.P. and K.H. wrote the manuscript.

## 8 Funding

This study was supported by Marsden Fund grant U001617 (K.H.) from the Royal Society of New Zealand Te Apārangi.

## 9 Acknowledgments

We would like to thank Kunling Li for his help in collecting behavioral data.

## 10 Data Availability

Datasets are available on request. The raw data supporting the conclusions of this manuscript will be made available by the authors, without undue reservation, to any qualified researcher.

## Notes

https://github.com/blakeporterneuro/weightLiftingTask

## References

Arac, A., P. Zhao, B. H. Dobkin, S. T. Carmichael and P. Golshani (2019). DeepBehavior: A Deep Learning Toolbox for Automated Analysis of Animal and Human Behavior Imaging Data. Front Syst Neurosci, 13, 20.

Bardgett, M. E., M. Depenbrock, N. Downs, M. Points and L. Green (2009). Dopamine modulates effort-based decision making in rats. Behav Neurosci, 123(2), 242–251.

Cowen, S. L., G. A. Davis and D. A. Nitz (2012). Anterior cingulate neurons in the rat map anticipated effort and reward to their associated action sequences. J Neurophysiol, 107(9), 2393–2407.

Floresco, S. B. and S. Ghods-Sharifi (2007). Amygdala-prefrontal cortical circuitry regulates effort-based decision making. Cereb Cortex, 17(2), 251–260.

Floresco, S. B., M. T. Tse and S. Ghods-Sharifi (2008). Dopaminergic and glutamatergic regulation of effort- and delay-based decision making. Neuropsychopharmacology, 33(8), 1966–1979.

Harris, A. and S. L. Lim (2016). Temporal Dynamics of Sensorimotor Networks in Effort-Based Cost-Benefit Valuation: Early Emergence and Late Net Value Integration. J Neurosci, 36(27), 7167–7183.

Hart, E. E., J. O. Gerson, Y. Zoken, M. Garcia and A. Izquierdo (2017). Anterior cingulate cortex supports effort allocation towards a qualitatively preferred option. Eur J Neurosci, 46(1), 1682–1688.

Hillman, K. L. and D. K. Bilkey (2010). Neurons in the rat anterior cingulate cortex dynamically encode cost-benefit in a spatial decision-making task. J Neurosci, 30(22), 7705–7713.

Holec, V., H. L. Pirot and D. R. Euston (2014). Not all effort is equal: the role of the anterior cingulate cortex in different forms of effort-reward decisions. Front Behav Neurosci, 8, 12.

Karimi, S., A. Mesdaghinia, Z. Farzinpour, G. Hamidi and A. Haghparast (2017). Reversible inactivation of the lateral hypothalamus reversed high reward choices in cost-benefit decision-making in rats. Neurobiol Learn Mem, 145, 135–142.

Kennerley, S. W., A. F. Dahmubed, A. H. Lara and J. D. Wallis (2009). Neurons in the frontal lobe encode the value of multiple decision variables. J Cogn Neurosci, 21(6), 1162–1178.

Klein-Flugge, M. C., S. W. Kennerley, K. Friston and S. Bestmann (2016). Neural Signatures of Value Comparison in Human Cingulate Cortex during Decisions Requiring an Effort-Reward Trade-off. J Neurosci, 36(39), 10002–10015.

Krakauer, J. W., A. A. Ghazanfar, A. Gomez-Marin, M. A. MacIver and D. Poeppel (2017). Neuroscience Needs Behavior: Correcting a Reductionist Bias. Neuron, 93(3), 480–490.

Kurniawan, I. T., B. Seymour, D. Talmi, W. Yoshida, N. Chater and R. J. Dolan (2010). Choosing to make an effort: the role of striatum in signaling physical effort of a chosen action. J Neurophysiol, 104(1), 313–321.

Lindenbach, D., J. K. Seamans and A. G. Phillips (2019). Activation of the ventral subiculum reinvigorates behavior after failure to achieve a goal: Implications for dopaminergic modulation of motivational processes. Behav Brain Res, 356, 266–270.

Ma, L., J. M. Hyman, A. G. Phillips and J. K. Seamans (2014). Tracking progress toward a goal in corticostriatal ensembles. J Neurosci, 34(6), 2244–2253.

Mathis, A., P. Mamidanna, K. M. Cury, T. Abe, V. N. Murthy, M. W. Mathis and M. Bethge (2018). DeepLabCut: markerless pose estimation of user-defined body parts with deep learning. Nat Neurosci, 21(9), 1281–1289.

Pessiglione, M., L. Schmidt, B. Draganski, R. Kalisch, H. Lau, R. J. Dolan and C. D. Frith (2007). How the brain translates money into force: a neuroimaging study of subliminal motivation. Science, 316(5826), 904–906.

Porter, B. S., K. L. Hillman and D. K. Bilkey (2019). Anterior cingulate cortex encoding of effortful behavior. J Neurophysiol, 121(2), 701–714.

Proulx, C. D., S. Aronson, D. Milivojevic, C. Molina, A. Loi, B. Monk, S. J. Shabel and R. Malinow (2018). A neural pathway controlling motivation to exert effort. Proc Natl Acad Sci U S A, 115(22), 5792–5797.

Randall, P. A., C. A. Lee, S. J. Podurgiel, E. Hart, S. E. Yohn, M. Jones, M. Rowland, L. Lopez-Cruz, M. Correa and J. D. Salamone (2014). Bupropion increases selection of high effort activity in rats tested on a progressive ratio/chow feeding choice procedure: implications for treatment of effort-related motivational symptoms. Int J Neuropsychopharmacol, 18(2).

Randall, P. A., M. Pardo, E. J. Nunes, L. Lopez Cruz, V. K. Vemuri, A. Makriyannis, Y. Baqi, C. E. Muller, M. Correa and J. D. Salamone (2012). Dopaminergic modulation of effort-related choice behavior as assessed by a progressive ratio chow feeding choice task: pharmacological studies and the role of individual differences. PLoS One, 7(10), e47934.

Robinson, M. J., S. M. Warlow and K. C. Berridge (2014). Optogenetic excitation of central amygdala amplifies and narrows incentive motivation to pursue one reward above another. J Neurosci, 34(50), 16567–16580.

Rudebeck, P. H., M. E. Walton, A. N. Smyth, D. M. Bannerman and M. F. Rushworth (2006). Separate neural pathways process different decision costs. Nat Neurosci, 9(9), 1161–1168.

Salamone, J. D., M. N. Arizzi, M. D. Sandoval, K. M. Cervone and J. E. Aberman (2002). Dopamine antagonists alter response allocation but do not suppress appetite for food in rats: contrast between the effects of SKF 83566, raclopride, and fenfluramine on a concurrent choice task. Psychopharmacology (Berl), 160(4), 371–380.

Salamone, J. D., M. Correa, S. Ferrigno, J. H. Yang, R. A. Rotolo and R. E. Presby (2018). The Psychopharmacology of Effort-Related Decision Making: Dopamine, Adenosine, and Insights into the Neurochemistry of Motivation. Pharmacol Rev, 70(4), 747–762.

Salamone, J. D., M. S. Cousins and S. Bucher (1994). Anhedonia or anergia? Effects of haloperidol and nucleus accumbens dopamine depletion on instrumental response selection in a T-maze cost/benefit procedure. Behav Brain Res, 65(2), 221–229.

Schweimer, J. and W. Hauber (2005). Involvement of the rat anterior cingulate cortex in control of instrumental responses guided by reward expectancy. Learn Mem, 12(3), 334–342.

Schweimer, J. and W. Hauber (2006). Dopamine D1 receptors in the anterior cingulate cortex regulate effort-based decision making. Learn Mem, 13(6), 777–782.

Treadway, M. T., N. A. Bossaller, R. C. Shelton and D. H. Zald (2012). Effort-based decision-making in major depressive disorder: a translational model of motivational anhedonia. J Abnorm Psychol, 121(3), 553–558.

Varazzani, C., A. San-Galli, S. Gilardeau and S. Bouret (2015). Noradrenaline and dopamine neurons in the reward/effort trade-off: a direct electrophysiological comparison in behaving monkeys. J Neurosci, 35(20), 7866–7877.

Walton, M. E., D. M. Bannerman and M. F. Rushworth (2002). The role of rat medial frontal cortex in effort-based decision making. J Neurosci, 22(24), 10996–11003.

Yang, X. H., J. Huang, C. Y. Zhu, Y. F. Wang, E. F. Cheung, R. C. Chan and G. R. Xie (2014). Motivational deficits in effort-based decision making in individuals with subsyndromal depression, first-episode and remitted depression patients. Psychiatry Res, 220(3), 874–882.

