## Supplemental Data for "A novel weight lifting task for investigating effort and persistence in rats"

Supplementary Material

### 1 Supplementary Figures

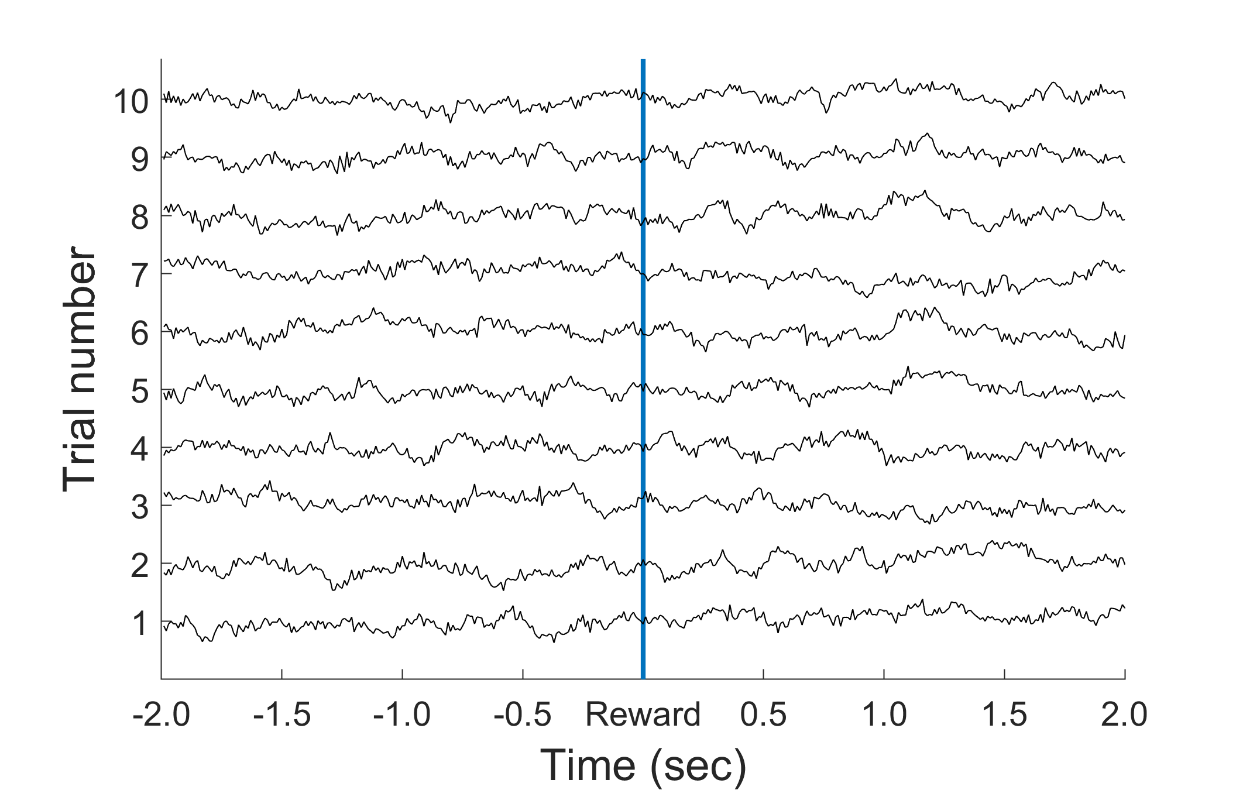

**Supplemental Figure 1:** Example local field potential traces from the anterior cingulate cortex in a rat carrying out the WLT. Data are shown for 10 successful trials on 135 g centered on the time the rat triggers the reward (blue line), i.e. the weight has reached 30 cm off the base and triggered the reed switch. Local field potential traces remain stable and free of artefacts while the rat pulls the rope (the 2 seconds prior to the reward time) then drops the rope and moves to the reward spout (the 2 seconds after the reward time).

### 2 Supplementary Videos

**Supplemental Video 1:** WLT training process showing rats advance from stages i through iv.

**Supplemental Video 2:** A rat performing a trial of the WLT on 0 g.
